# Cellomics approach for high-throughput functional annotation of *Caenorhabditis elegans* neural network

**DOI:** 10.1101/182923

**Authors:** Wataru Aoki, Hidenori Matsukura, Yuji Yamauchi, Haruki Yokoyama, Koichi Hasegawa, Ryoji Shinya, Mitsuyoshi Ueda

## Abstract

Even in *Caenorhabditis elegans*, which has only 302 neurons, relationships between behaviors and neural networks are not easily elucidated. In this study, we proposed a novel cellomics approach enabling high-throughput and comprehensive exploration of the functions of a single neuron or a subset of neurons in a complex neural network on a particular behavior. To realize this, we combined optogenetics and Brainbow technologies. Using these technologies, we established a *C. elegans* library where opsin is labeled in a randomized pattern. Behavioral analysis on this library under light illumination enabled high-throughput annotation of neurons affecting target behaviors. We applied this approach to the egg-laying behavior of *C. elegans* and succeeded in high-throughput confirmation that hermaphrodite-specific neurons play an important role in the egg-laying behavior. This cellomics approach will lead to the accumulation of neurophysiological and behavioral data of the *C. elegans* neural network, which is necessary for constructing neuroanatomically grounded models of behavior.

## INTRODUCTION

One of the primary objectives of neuroscience is to understand how computations are implemented across neural networks. However, neural networks are so complex that it is not easy to elucidate how they give rise to certain behavior. Hermaphrodites of *Caenorhabditis elegans*, for instance, have a simple neural network consisting of 302 neurons, and their connectome, a comprehensive map of neural connections, has already been clarified^1,2^. However, even in *C. elegans*, the relationships between behaviors and neural networks have not been comprehensively revealed.

Data-driven science, in which an enormous amount of data is accumulated in a hypothesis-free manner, has deepened our understanding of complex biological processes. For example, the applications of state-of-the-art mass spectrometry^3^ and next-generation sequencers^4^ have made it easy to acquire omics data on various biological strata, including those concerning genome, transcriptome, proteome, and metabolome, thus streamlining the process of understanding diverse biological processes. However, even with such omics analyses, it is still difficult to gain deeper insights into individual-level biological processes engendered by complex neural networks.

To understand such biological processes derived from complex cellular networks, it is important to develop a new methodology for collecting omics data at the cellular level (cellomics) and implement data-driven science at the individual-level. In recent studies, an approach to reveal cellular network structures comprehensively, which can be called “descriptive” cellomics, has been gradually changing the landscape of neuroscience. Serial electron microscopy^5^, for example, has been proven to be useful in determining connectomes in various model organisms and promoting comparative connectomics in *C. elegans*^6,7^. Moreover, researchers have also developed new approaches, such as Brainbow^8^ or Optobow^9^, which involve sparse labeling of fluorescent proteins or opsins in an attempt to reveal brain structures using optical microscopy. Recent improvements in brain visualization^10,11^ and light-sheet microscopy^12^ technologies have made it possible to perform high-speed imaging of the mammalian whole brain. These studies have already benefited the field of neuroscience through the accumulation of data on the structure of the brain in a hypothesis-free manner.

To understand the workings of the brain, its structural information is important, but that alone is still insufficient. It is considered essential to accumulate comprehensively neurophysiological and behavioral data on what functions a particular part of the brain has, in an effort to construct neuroanatomically grounded models of behavior. In this context, we considered establishing a novel methodology called “functional” cellomics, which allows comprehensive exploration of what functions a single neuron or a subset of neurons in a complex neural network has on a certain behavior. This sort of approach has already been demonstrated for human brains in the form of natural experiments effectuated by diseases, accidents, wars, and other incidents. Throughout history, loss of function and gain of function screening by natural experiments has caused various serendipitous discoveries, such as the functions of the hippocampus^13^, which have formed the basis of modern neuroscience. If similar experiments can be performed in a high-throughput and systematic manner at the neural circuit level, they will provide a lot of knowledge in the field of neuroscience.

In this study, we attempted to demonstrate that functional cellomics is feasible using *C. elegans* and is a promising approach for the comprehensive functional annotation of neural networks. A methodology to realize functional cellomics is required to allow researchers to manipulate a neuron or a subset of neurons in a high-throughput, hypothesis-free, single-cell-resolution, and simple manner and to quantify the effect of such manipulation on a behavior. In *C. elegans* research, various methodologies for collecting neurophysiological data are available, such as calcium imaging^14^, optical and electrophysiological recording^15^, and laser ablation^16^. However, no single methodology satisfies all of the criteria necessary for realizing the conceptual framework of functional cellomics.

To overcome this obstacle and achieve this approach, we fused optogenetics and Brainbow technologies. Optogenetics is a technique that enables on-demand photoregulation of neural activity through the expression of opsin genes^17^. In the field of optogenetics, a wide array of tools for activating, suppressing, and killing neurons has been developed and changed the methodology of neuroscience. However, with conventional optogenetics, it is essential to determine in advance which promoters should be used to produce an opsin in specific neurons^18^. Therefore, this approach is hypothesis-driven, meaning that it is effective in precisely testing existing hypotheses, but is not conducive to establishing entirely new ones (Figure 1a).

**Figure 1.**
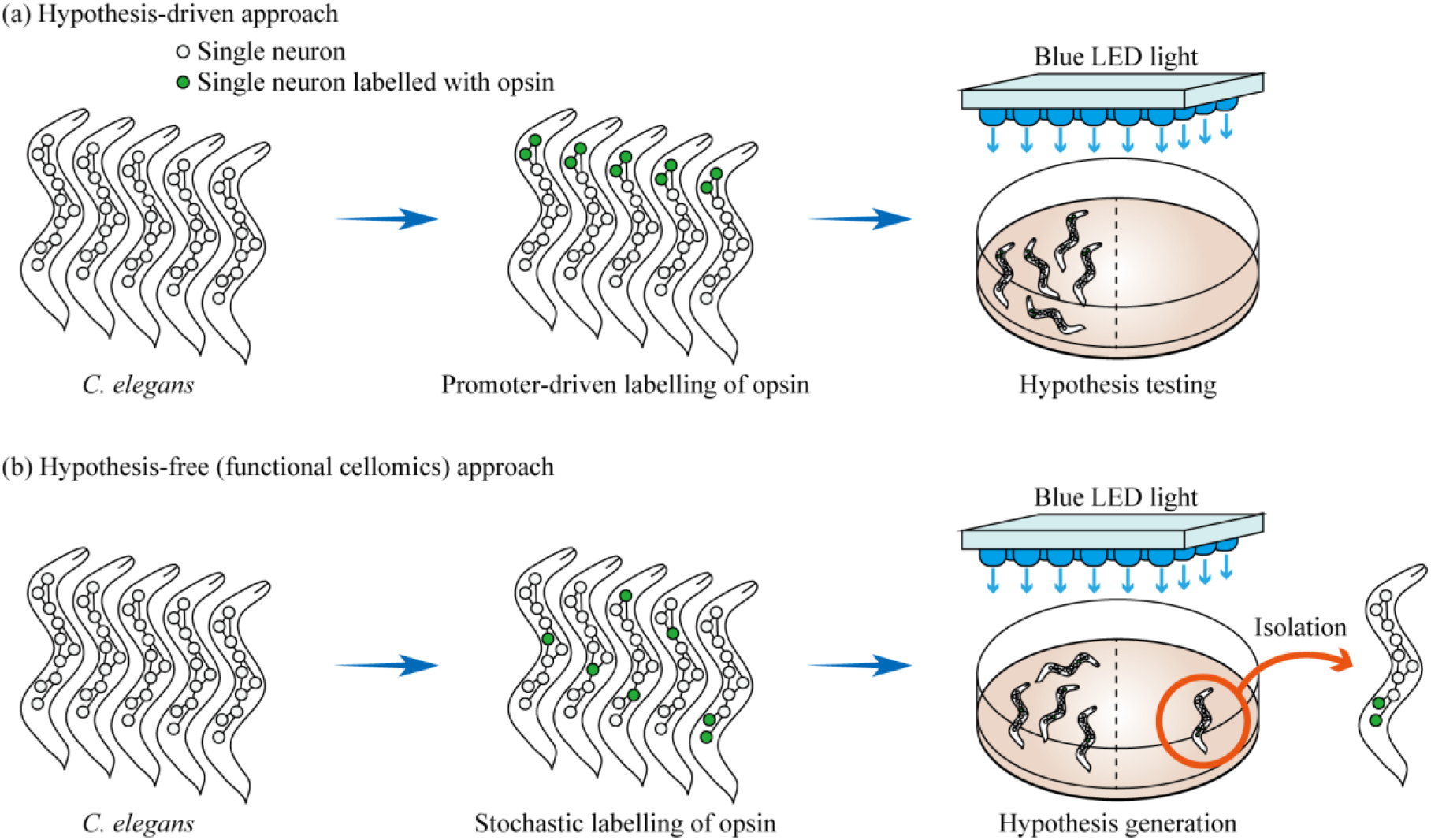
Comparison between hypothesis-driven and hypothesis-free (functional cellomics) approaches for annotation of neural networks. (a) Experimental scheme for hypothesis-driven neural network analysis. First, a hypothesis is formulated as to which neuron or a subset of neurons is responsible for a certain behavior. Next, only specific neurons are allowed to express an effector (opsin in this figure) by using appropriate promoters. Finally, behavioral experiments are conducted with this transgenic *C. elegans* to test whether the initial hypothesis is correct. (b) Experimental scheme for hypothesis-free (functional cellomics) approach for neural network analysis. In the method that we proposed, it is determined in a stochastic manner whether or not each single neuron is labeled by an effector. By performing behavioral experiments on a *C. elegans* library in which neurons are labeled in various patterns, one can expect to find individuals that differ behaviorally from wild-type *C. elegans*. By isolating these individuals and examining which neuron was producing opsin, one may be able to discover a new relationship between neural networks and behavior.

To address this issue, we designed a new optogenetic experimental scheme to accomplish hypothesis-free neural network annotation (Figure 1b). Specifically, we attempted to develop a system that stochastically determines whether the effector is produced in each single neuron and to acquire a *C. elegans* library in which the effector is labeled in diverse patterns. To achieve this stochastic labeling, we adopted Brainbow technologies based on the Cre-lox system^8^. If a *C. elegans* library with a random labeling pattern of the effector can be obtained through the use of Brainbow technologies, it would be possible to uncover hitherto unknown relationships between neural networks and behaviors in a high-throughput manner by performing behavioral experiments with the library under light illumination. This approach is hypothesis-free because individuals showing abnormal behavior are first detected and responsible neurons are then identified. By analogy, functional cellomics can be regarded as a cell level application of the forward genetics concept, in which random mutations are introduced across the whole genome and genes altered in mutants showing some phenotypic changes are then identified.

In the present study, we successfully demonstrated the possibility that neurons affecting a behavior of interest can be identified by the combined use of optogenetics and Brainbow technologies. The results substantiated the basic concept of functional cellomics that enables functional annotation of neural networks in a high-throughput, hypothesis-free, single-cell-resolution, and simple manner.

## RESULTS

### Design of strategy for optogenetic functional cellomics

A methodology to achieve functional cellomics is required to accomplish stochastic labeling of individual neurons with an effector gene in a high-throughput, hypothesis-free, single-cell-resolution, and simple manner. Then, we focused on Brainbow technologies^8^. Brainbow technologies refer to systems that can stochastically determine whether a certain gene is expressed in a certain cell through the application of the Cre-lox recombination system. In Brainbow technologies, multiple *lox* variants (e.g., *loxP* and *lox2272* sequences) are inserted alternately downstream of one promoter and two other genes are interposed between these *lox* sequences (Figure 2). If Cre recombinase is allowed to act on this sequence, excision occurs exclusively either between *loxP* sequences or between *lox2272* sequences. Consequently, it becomes possible to determine which of these two genes is expressed in a Cre-dependent manner.

**Figure 2.**
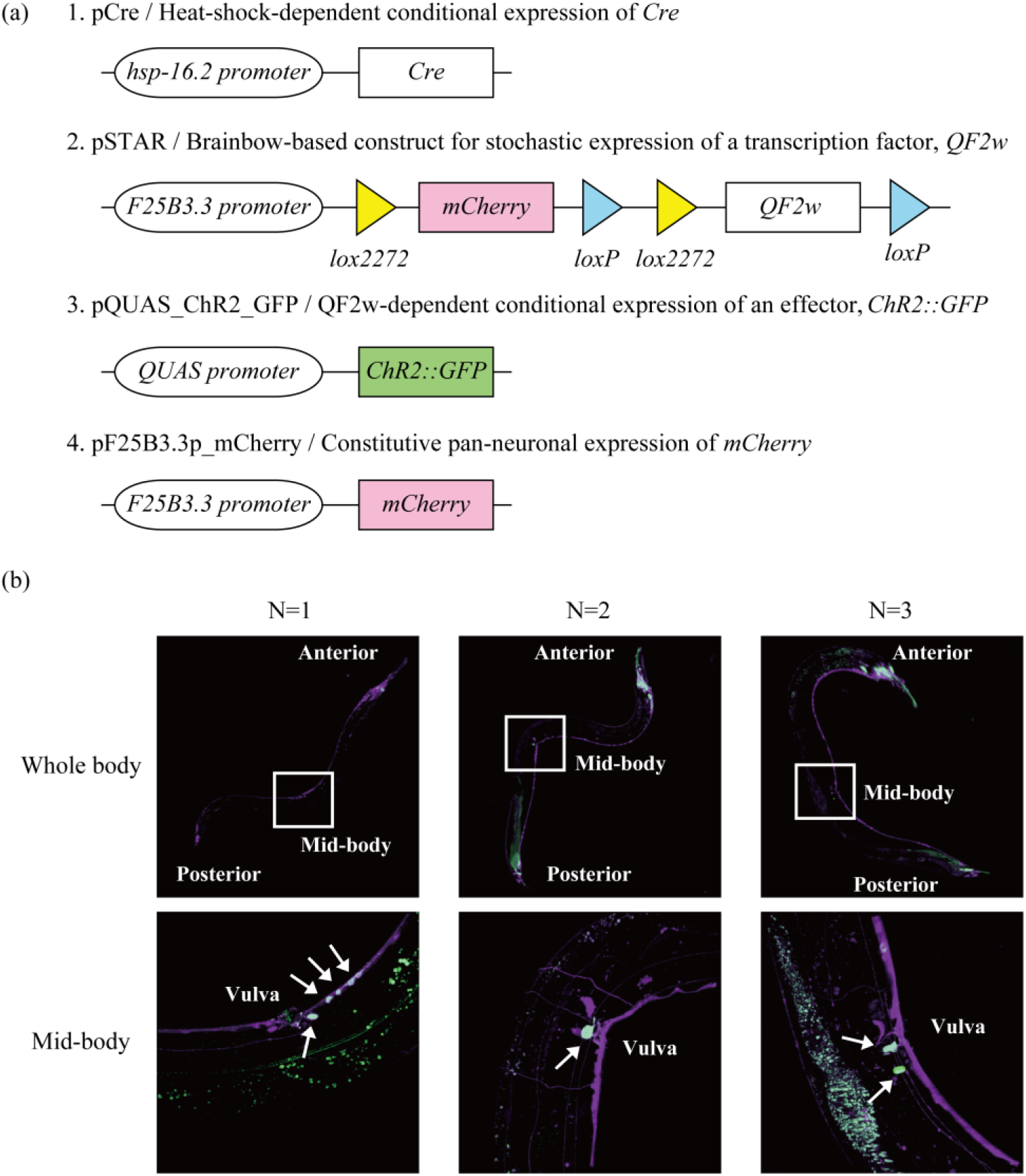
Stochastic labeling of neurons based on Brainbow technologies. (a) One of the constructs (pCre) is designed to produce Cre recombinase in a heat-shock-dependent manner. Another construct (pSTAR) is based on Brainbow technologies, with the *lox* sequences, *mCherry*, and the transcription factor *QF2*^*w*^ encoded downstream of the pan-neuronal promoter (*F25B3.3p*). In the initial state, pSTAR produces mCherry in all neurons. When an excision by Cre occurs between *loxP* sequences, mCherry continues to be produced. When an excision occurs between *lox2272* sequences, on the other hand, QF2^w^ is produced. pQUAS_ChR2_GFP expresses the effector *ChR2::GFP* in a QF2^w^-dependent manner. pF25B3.3p_mCherry encodes *mCherry* downstream of the pan-neuronal promoter *F25B3.3p*. (b) Stochastic labeling of neurons with ChR2-GFP. A brief heat shock was applied to the transgenic *C. elegans* that carries the four constructs as extrachromosomal (Ex) arrays, and each sample was observed using confocal laser scanning microscopy 12 h later. The whole body was observed at a magnification of 10×, and the mid-body section (framed in white in the whole body picture), where neurons can be counted with relative ease, was observed at a magnification of 40×. The 2D images shown are the maximum-intensity projection reconstructed from the z-stacks of the images acquired with the confocal microscope. The fluorescence of ChR2-GFP is presented in green, and that of mCherry in magenta. The cells expressing both fluorescent proteins are presented in white and indicated by an arrow. The white-dyed cells differed from one individual to another, indicating the success of stochastic labeling.

To implement functional cellomics, we designed four plasmids (Figure 2a). The plasmid pCre expresses Cre recombinase in response to a heat shock. In the plasmid pSTAR, *lox* sequences, *mCherry*, and a transcription factor (*QF2*^*w* 19^) were inserted downstream of a pan-neuronal promoter (*F25B3.3p*). The plasmid pQUAS_ChR2_GFP expresses an arbitrary effector gene in a QF2^w^-dependent manner. In the present study, we adopted channelrhodopsin-2 fused with GFP (*ChR2::GFP*). Since the constructs producing a transcription factor or an effector are modularized, it is easy to use not only opsin but also various other effectors. We also constructed pF25B3.3p_mCherry, which continuously expresses mCherry even after Cre recombination.

When all of these plasmids are introduced into *C. elegans*, all neurons produce only mCherry at the initial state. After a heat shock is applied to induce Cre recombinase, the production of mCherry continues if the genome is excised between *loxP* sequences, while QF2^w^ is instead produced if an excision occurs between *lox2272* sequences. In neurons producing QF2^w^, ChR2-GFP is produced as an effector, enabling the on-demand activation of these neurons by light illumination. Since GFP is fused to ChR2, it is easy to identify which neurons are producing opsin following a behavioral experiment.

### Stochastic labeling of neurons at single-cell level

We introduced the above-mentioned four plasmids into *C. elegans* to establish the AYK338 strain (*aykEx338 [hsp-16.2p::Cre, F25B3.3p::lox2272::mCherry::loxP::lox2272::QF2*^*w*^*::loxP, QUAS::ChR2::GFP, F25B3.3p:: mCherry]*). After this *C. elegans* strain had propagated, a brief heat shock was applied to determine whether ChR2-GFP was labeled stochastically in each *C. elegans* individual. After isolating three individuals, we observed their mid-body sections where the neuron density was low at a magnification of 40×. The results revealed that the labeling pattern of ChR2-GFP differed from one individual to another (Figure 2b).

### Identification of neurons responsible for egg-laying behavior

In functional cellomics, stochastic labeling of an effector gene makes it possible to explore the relationships between neural networks and behaviors in a hypothesis-free and comprehensive manner. To demonstrate the feasibility of functional cellomics, we selected the egg-laying behavior of *C. elegans* as a model. It is known that a relatively simple neural network is responsible for controlling the egg-laying behavior of this nematode. In this behavior, two hermaphrodite-specific neurons (HSNs: HSNR and HSNL) play a central role through direct excitation of vulval muscles and ventral C neurons (VCs)^20–22^. Moreover, it is known that the activation of the HSNs by ChR2 induces egg-laying behavior^23^. If the HSNs can be identified by functional cellomics in a high-throughput manner, it shows that this strategy actually works.

We constructed a *C. elegans* library in which ChR2-GFP is stochastically labeled. Individuals of this transgenic *C. elegans* were filmed for 30 sec while illuminated with blue light (Figure 3a). Among the filmed individuals, 65% laid eggs in a light-dependent manner, whereas 35% did not (Figure 3bc). When a similar experiment was conducted without all-*trans* retinal (ATR), a cofactor of ChR2, no egg-laying behavior was observed. These results indicate that the egg-laying behavior observed in this experiment is ChR2-dependent, and that individual nematodes exhibiting the target phenotype can be readily obtained through the stochastic labeling of an effector.

**Figure 3.**
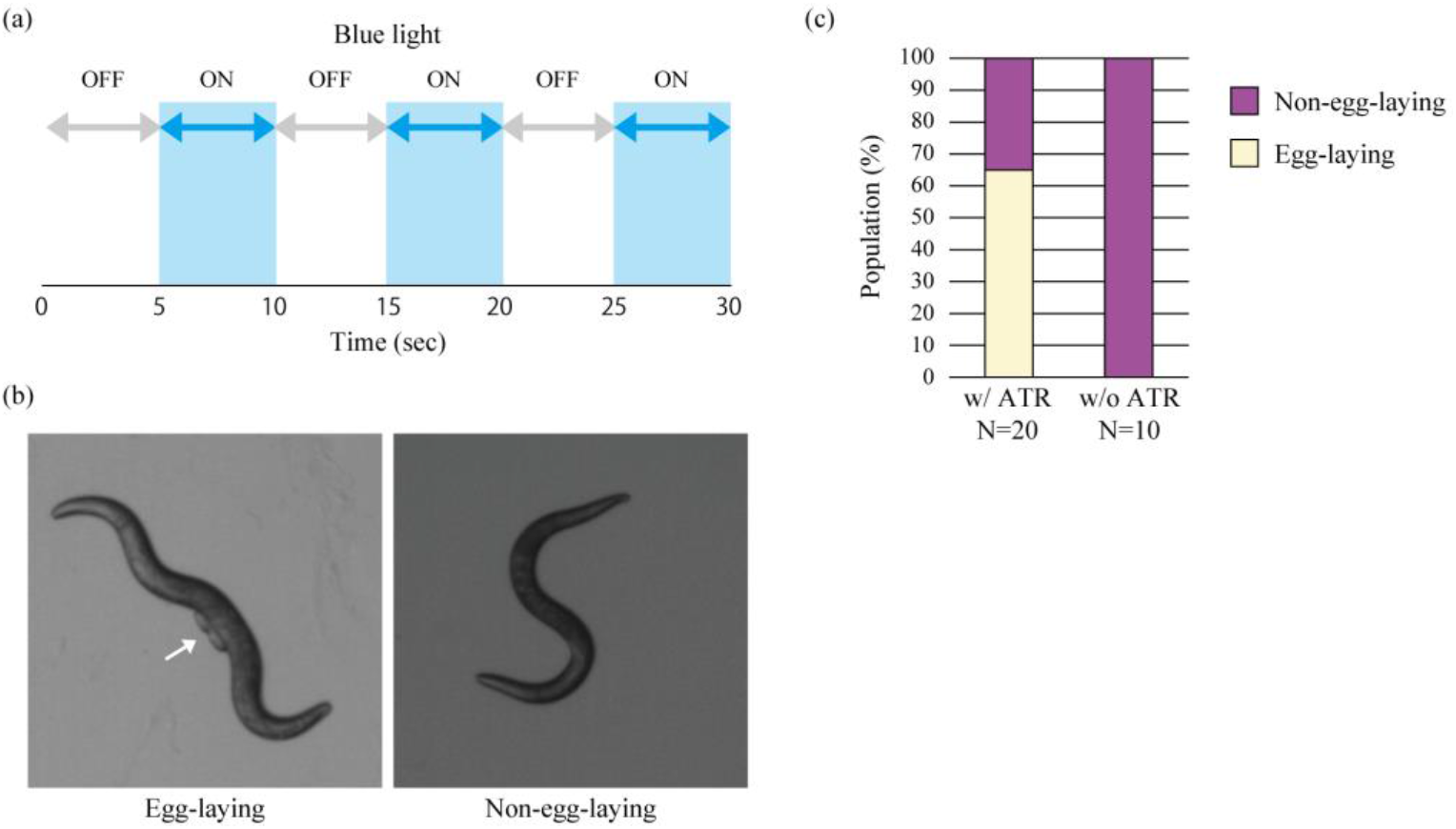
Acquisition of individuals exhibiting egg-laying behavior in a light-dependent manner. (a) Outline of the optogenetics experiment. The behavior of each nematode was filmed for 30 sec. During the filming, blue light was turned on and off at 5-second intervals. (b) Images captured from the representative footage during the behavioral experiment, showing an individual that exhibited egg-laying behavior in a light-dependent manner (left) and an individual that did not (right). The arrow indicates where the eggs are. (c) Proportion of individuals that exhibited light-dependent egg-laying behavior. In the presence of all-*trans* retinal (ATR), a cofactor of opsin, 65% of adult individuals exhibited egg-laying behavior under blue light. A negative control experiment was performed identically without ATR, in which all adult individuals turned out to be non-egg-laying.

Next, we used confocal laser scanning microscopy to determine whether ChR2-GFP was produced in the HSNs. After isolating egg-laying and non-egg-laying individuals and observing the vicinity of the vulva, we confirmed that GhR2-GFP was produced in the HSNs in the egg-laying individuals (Figure 4a), but it was not in the non-egg-laying individuals (Figure 4b). In the representative individual shown in Figure 4a, the production of ChR2-GFP was seen in HSNR but not in HSNL. A previous study demonstrated that killing one HSN by laser ablation did not markedly affect the egg-laying behavior of the nematode, whereas killing both HSNs resulted in strong inhibition of the egg-laying behavior^21^. Our result that the egg-laying behavior was induced sufficiently by activating only one HSN is consistent with that of this previous study.

**Figure 4.**
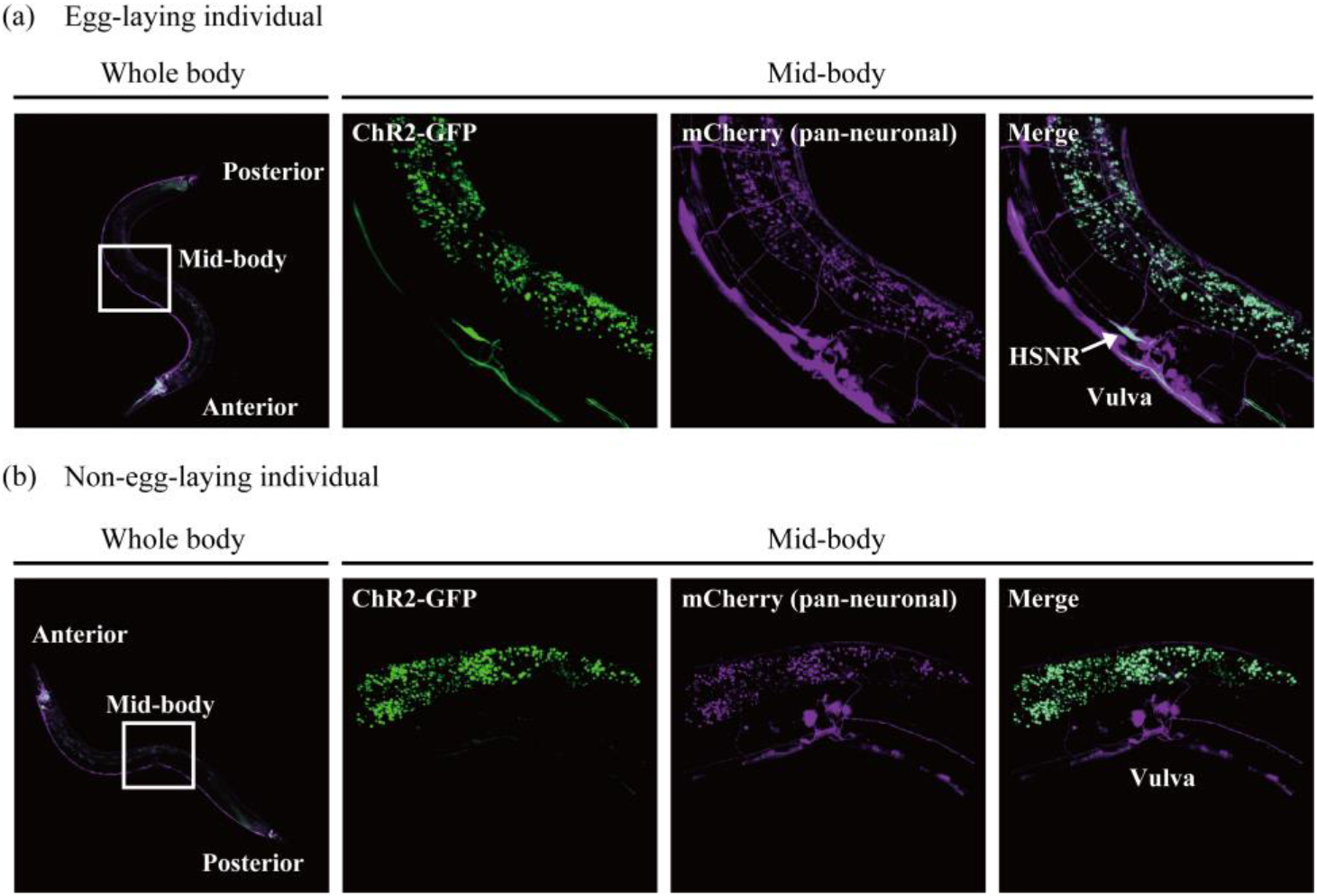
Identification of HSNs. The whole body was observed with a 10× objective, and the mid-body section with a 40× objective. The 2D images shown are the maximum-intensity projection reconstructed from the z-stacks of the images acquired with a confocal microscope. (a) Fluorescence images of an egg-laying individual. The fluorescence of ChR2-GFP is presented in green, and that of mCherry in magenta. The cells that are producing both fluorescent proteins are presented in white. In this particular individual that laid eggs in a light-dependent manner, the expression of ChR2-GFP in HSNR was observed. (b) Fluorescence image of a non-egg-laying individual. Light did not induce egg-laying behavior in this individual, and no production of ChR2-GFP was observed in neurons around the vulva.

## DISCUSSION

Although various methodologies have been established to explore the properties of neural networks, no single methodology satisfies all of the criteria necessary for realizing the conceptual framework of functional cellomics. One typical example of existing methods, which is very easy yet effective, is to induce the production of effectors using cell-type-specific promoters. However, this approach is basically hypothesis-driven because one needs to select specific promoters a priori, meaning that it is not suitable for establishing entirely new hypotheses. Moreover, it is also difficult to analyze neural networks at single-cell resolution because *C. elegans* has few single-cell-specific promoters. By extension, data-driven neuron-behavior mapping is not impracticable with this approach. In *Drosophila melanogaster*, neuron-behavior mapping was attempted using 1049 distinct cell-type-specific GAL4 lines to selectively target ChR2^24^. As a result, 29 discrete, statistically distinguishable behavioral phenotypes were discovered, indicating that data-driven neuron-behavior mapping is a useful approach. However, owing to the fact that a large number of *D. melanogaster* lines are required, this method is costly and laborious. Furthermore, as this method utilizes cell-type-specific promoters, neurons cannot be manipulated at single-cell resolution. Another example is laser ablation, which enables hypothesis-free and single-cell-resolution analysis^16^. Although this is a powerful technique applicable to any species, its low throughput makes it difficult to conduct experiments involving various patterns of intervention. Additionally, laser ablation lacks expandability in that it cannot activate or suppress neurons. A recent study has suggested the possibility that contradictory results are generated depending on the mode of intervention^25^, indicating the need to compare results obtained from various modes of intervention (activation, suppression, killing, etc.). Another method enabling analysis at single-cell resolution involves application of a heat shock to only specific cells to induce effectors^26,27^. This method is capable of manipulating neurons in many ways; however, similar to laser ablation, it is also restricted by its low throughput capacity. The patterned illumination technique using digital micromirror devices, whose development has been advancing in recent years, allows for a high degree of freedom in experimental design and has a relatively high throughput^23^. However, it is still difficult to perform accurate analysis at single-cell resolution with this method because multiple adjacent neurons may be illuminated simultaneously, unless sufficiently sparse expression patterns of effectors are provided^28^.

Functional cellomics described in this study is the first approach to combine all properties necessary for achieving individual-level cellomics, that is, high-throughput, hypothesis-free, single-cell resolution, and simplicity. In fact, by applying functional cellomics to the egg-laying behavior of *C. elegans*, we managed to establish the proof of concept of this approach.

Compared with the existing methodologies, this system has advantages in terms of throughput, resolution, and expandability. First, it can be easily implemented in any laboratory without requiring any specialized equipment. Second, having no limitations in feasible labeling patterns, it is completely hypothesis-free, facilitating easy labeling at singlecell resolution, even for bilaterally symmetrical neuron pairs with almost identical gene expression patterns. Third, with one transgenic *C. elegans* individual capable of propagating many other individuals with different labeling patterns, it is both simple and high-throughput. Fourth, it can intervene in neural networks in various fashions. Besides opsin, which was employed in the present study, any effectors can be used as long as they cause either loss of function or gain of function in neurons. This enables a variety of interventions, such as cell killing^29,30^, suppression^31^, activation^32^, and gene expression control^33,34^. Fifth, multitudes of experimental designs are available using *C. elegans* promoterome^35^ and bipartite gene expression systems such as QF2^w^ and Gal4^36^. For example, instead of a pan-neuronal promoter, one can use a promoter specific to a subset of neurons for more “focused” functional cellomics. In addition, simultaneously employing QF2^w^ and Gal4, one can label multiple effectors in a stochastic fashion.

Although promising, there are two points of concern regarding functional cellomics. One is the high probability of effector labeling, which makes it difficult to calculate the labeling rate accurately. To perform a well-designed experiment, it is necessary to achieve strict control of the probability of effector labeling, as is the case with forward genetics in which the mutation rate is predetermined. Figure 3c shows that a large population of *C. elegans* (65%) displayed egg-laying behavior, and this was probably because a large proportion of neurons were labeled by opsin. When too many neurons are labeled by effectors, it becomes difficult to determine which neurons are responsible for the target behavior. Concomitant use of single-copy integration and *lox* variants may be able to control the probability of labeling only a desired number of all 302 neurons of a *C. elegans* hermaphrodite. In our laboratory, we are proceeding with the construction of a more sophisticated system that will help improve this issue.

The second point is how to ensure that the obtained results are reproducible. If functional cellomics implies that a certain labeling pattern may affect the target behavior, it is still necessary to verify this by other methods. To reproduce the labeling pattern, methodologies that can evoke gene expression in arbitrary cells, such as use of a pulsed infrared laser^26^ and multi-step optogenetics^27^, may be applied to verify the results relatively easily. Besides, the intersectional Cre-lox strategy^37^ and multiple-feature Boolean logic^38^ may also be applicable to reproducing the labeling pattern.

In conclusion, we have demonstrated for the first time the possibility of identifying neurons responsible for a target behavior by randomizing the labeling patterns of effector genes based on Brainbow technologies. The results substantiated the basic concept of functional cellomics, which enables functional annotation of neural networks of *C. elegans* in a high-throughput, hypothesis-free, single-cell-resolution, and simple manner. Since its connectome information is already mapped and available, *C. elegans* is highly compatible with functional cellomics. By overlaying the connectome information with the results of cyclopedic intervention experiments by functional cellomics, we should become able to construct neuroanatomically grounded models of behavior that can explain how complex neural networks implement computation.

## METHODS

### Construction of plasmids

To construct pCre, *SV40NLS-Cre* was amplified from pPGK-Cre-bpA deposited by Klaus Rajewsky (Addgene plasmid #11543). The amplified fragment was inserted into pPD49_78 deposited by Andrew Fire (Addgene plasmid #1447).

To construct pSTAR, a backbone plasmid with *lox* and *QF2*^*w* 19^ sequences was synthesized (Thermo Fisher Scientific, MA, USA). In addition, *mCherry* was subcloned from pGH8^39^ deposited by Erik Jorgensen (Addgene plasmid #19359) and a pan-neuronal promoter, *F25B3.3p*, was cloned from the *C. elegans* genome. These two fragments were inserted into the backbone plasmid.

To construct pQUAS_ChR2_GFP, *ChR2 (H134R)* and *GFP (S65C)* were amplified from pAAV-Ef1a-vCreDIO hChR2(H134R)-EYFP deposited by Karl Deisseroth (Addgene plasmid #55643) and L2680 deposited by Andrew Fire (Addgene plasmid #1516), respectively. In addition, a QF2^w^-dependent promoter sequence, *QUAS::Δpes-10*^40^, was synthesized (Thermo Fisher Scientific). These three fragments were inserted into pPD49_78.

To construct pF25B3.3p_mCherry, *F25B3.3p* and *mCherry* were subcloned from pSTAR. These fragments were inserted into pPD49_78.

### Culture conditions

Worms were cultivated on nematode growth medium (NGM) plates with *Escherichia coli* OP50. Specifically, the OP50 plates were prepared with 250 μL of OP50 seeded into 6cm NGM plates. The worms were maintained at 20°C, with care taken to ensure that the temperature shifted as little as possible. To perform optogenetic experiments, 300 μL of 500 μM ATR; Sigma-Aldrich, MO, USA) was added to solid NGM plates with *E. coli*, and the samples were allowed to dry while shielded from light by aluminum foil.

### Transgenic strains

Injections into the nematodes were performed with the aid of a stereomicroscope (SZX10; Olympus, Tokyo, Japan) equipped with a Femtojet 4i (5252 000.021; Eppendorf, Hamburg, Germany) and Femtotips II (1501040; Eppendorf). The strain AYK338 (*aykEx338 [hsp-16.2p::Cre, F25B3.3p::lox2272::mCherry::loxP::lox2272::QF2w::loxP, QUAS::ChR2::GFP, F25B3.3p:: mCherry]*) was generated by co-injecting the four plasmids constructed in this study (50 ng μL^-1^ each in water) into the *C. elegans* N2 background. The injection was performed on 20 nematodes with an N2 background, and three mCherry-expressing lines were obtained.

### Induction of Cre recombinase for stochastic labeling of ChR2

Transgenic worms were placed on NGM plates with or without ATR. The worms were incubated at 37°C for 30 min for the induction of Cre recombinase by heat shock, after which they were placed in an incubator at 20°C. At 12 h after the heat shock, the worms were examined by egg-laying assay and/or fluorescence microscopy.

### Fluorescence microscopy

A 5% agarose pad (01149-05; Nacalai Tesque, Kyoto, Japan) was prepared, onto which 5 μL of 50 mM sodium azide (830011; Nacalai Tesque) was placed. *C. elegans* worms were picked up and placed onto the agarose pad with sodium azide, over which a cover glass was placed gently. Fluorescence was observed by confocal laser scanning microscopy (LSM700; Carl Zeiss, Oberkochen, Germany). Fluorescence of GFP and mCherry was observed using 488 nm and 561 nm lasers, respectively. Acquired images were processed using Zen Lite, Imaris, or ImageJ^41^ software.

### Egg-laying assay

The worms were observed under a stereomicroscope (SZX10; Olympus) equipped with a camera (HAS-L1; DITECT, Tokyo, Japan). The SZX10’s halogen lamp (410849; PHILIPS, Amsterdam, the Netherlands) was fitted with an optical filter (Asahi Spectra, Tokyo, Japan) that blocks wavelengths below 600 nm to prevent ChR2-GFP from being activated within the worms during observation. To activate ChR2-GFP, the worms were illuminated using blue light (LDL2-98X30BL2; CCS, Kyoto, Japan) powered by PD3-5024-4-PI (CCS). To prevent the LED’s blue light from being detected by the camera, the object lens was fitted with an optical filter (Asahi Spectra) to block wavelengths below 570 nm.

For an egg-laying assay to examine light-dependent behavior modulation, each worm was transferred to a 6-cm agar NGM plate without *E. coli* and filmed for 30 sec. During this filming, the blue light was turned on and off at 5-second intervals, and the individuals that exhibited egg-laying behavior during 30 sec were defined as egg-laying individuals. Irrespective of the absence or presence of *E. coli*, the activation of the HSNs induces egg-laying behavior^23^.

## ACKNOWLEDGMENTS

We thank Kenta Terai and Michiyuki Matsuda for valuable advice on fluorescence imaging. We thank Crimson Interactive Pvt. Ltd. for their assistance with manuscript editing. This work was supported by PREST, JST (grant No. JPMJPR16F1), JSPS KAKENHI (grant No. JP17K19452), and Kyoto University Live Imaging Center.

## AUTHOR CONTRIBUTIONS

W.A. conceived the project. W.A., H.M., Y.Y., and H.Y. performed experiments and data analysis. K.T., M.M., K.H., R.S., and M.U. provided advice on method development. The manuscript was prepared by W.A. and edited by all co-authors.

